# Mothbox and Mothbot: automated light trap and data processing system for scalable insect monitoring

**DOI:** 10.64898/2025.12.03.692171

**Authors:** Hubert A. Szczygieł, Brianna Johns, Bernat Fortet, Daisy H. Dent, Kit Quitmeyer, Andrew Quitmeyer

## Abstract

1. Insects represent the most diverse group of organisms on Earth, and comprise the majority of known species, yet they are seldom accounted for in large-scale biodiversity monitoring systems and conservation planning.
2. We have developed the *Mothbox* - an open source automated light trap that makes insect monitoring accessible to non-specialists and scalable for scientific and conservation purposes - and the *Mothbot* - a machine learning tool and user interface for processing data from automated light traps. The Mothbox is portable, durable, and low-cost, while Mothbot prioritizes human-in-the-loop data validation of computer vision outputs to ensure data quality and aid in developing custom image reference libraries.
3. Mothbox has been extensively tested in field conditions, with > 185 deployments, spanning > 450 nights and > 100 sampling locations. We present proof-of-concept studies that examine shifts in insect activity and richness across i. time intervals throughout the night at a single location, and ii. different habitat types, to assess whether these patterns can be captured through automated sampling with Mothbox hardware and Mothbot processing.
4. As the field of automated insect monitoring continues to develop, Mothbox provides an affordable entry point for a range of stakeholders. It enables entomologists to explore new research questions by tracking high temporal resolution changes in insect activity, as well as systematically monitoring insects across landscapes. Mothbox will help build global insect-monitoring capacity via autonomous sampling, improving our ability to detect biodiversity loss and guide effective conservation action.

## 1 Introduction

The majority of the world’s species are insects, and most insects are active at night (Wong & Didham, 2024). Insects are incredibly valuable ecologically and economically, and rapidly declining in many parts of the world (Basset & Lamarre, 2019). Specialisation makes insects particularly relevant biodiversity indicators; for example, lepidoptera larvae often feed on only one species or genus of plant (Dyer et al., 2007), and adults can similarly be adapted to pollinate single host plant species (Hahn & Brühl, 2016), thus, monitoring Lepidoptera can provide inferences about the plant community (Duran et al., 2022; Tyler, 2020). However, insects are rarely included in conservation planning, despite their great diversity, ecological significance, and evidence of global decline, because of the difficulty of conducting standardised large-scale insect monitoring (Chowdhury et al., 2023; Høye et al., 2021).

Traditional approaches to monitoring nocturnal insects require specialised knowledge and considerable time investment (Montgomery et al., 2021). Light-sheeting, i.e. attracting nocturnal insects to an illuminated sheet, is an effective method for visually assessing insect populations or targeting particular taxa. However, documenting every insect is often impractical and requires researchers to work through the night (Duran et al., 2022). Bucket traps, which attract and capture insects, do not require ongoing supervision, but are typically deployed for the purpose of lethal collection and require extensive manual sample processing once collected (Brehm & Axmacher, 2006). New technological approaches to biodiversity monitoring also face unique challenges when applied to insects. For example, genetic approaches (e.g. DNA metabarcoding) remain expensive, rely on reference libraries with limited coverage for many regions and taxa (Alvarado-Robledo et al., 2024), and are poorly suited to assessing abundance (Sickel et al., 2023), while satellite-based remote-sensing lacks the resolution for direct monitoring of insects (Rhodes et al., 2022). To bypass difficulties with these monitoring methods, the first automated light trap that used a programmed light source and computer vision for identification was developed in 2021 (Bjerge et al., 2021). This proof of concept inspired the creation of a range of automated insect monitoring systems, including the Automatic Moth Trap (Möglich et al., 2023), Automated Monitoring of Insects (Lord et al., 2026), DIOPSIS (Huijbers et al., 2025), and others (Beuchert, 2024; Hobern, 2021; Korsch et al., 2023). Automated non-lethal insect monitoring remains an emerging field, and existing devices, including the one presented in this manuscript, represent complementary approaches that balance cost, portability, image resolution, insect-attraction lighting, 4G connectivity, ease of modification, and durability as the research community collectively works toward further technological refinement.

## 2. Mothbox and Mothbot: An open-source automated light trap and image-processing platform

Here we describe the Mothbox, an automated light trap designed for scalable insect monitoring, and Mothbot, a platform for automated light trap data processing and validation. The Mothbox device integrates a low-cost Raspberry Pi computer, a high-resolution camera, and specialized attractant lights to capture images of nocturnal insects. The Mothbox has been developed specifically to optimize portability, adaptability, high-resolution images, and weather resistance.

Furthermore, the low materials cost of the latest model (Mothbox Pro; approximately USD 375) enables the simultaneous deployment of multiple units. Mothbot is an open-source tool designed to extract insect occurrence data from images captured by Mothbox and similar devices. We present two proof-of-concept studies from Panama that employ the Mothbox and Mothbot to detect and identify night-flying insects, demonstrating the potential for biodiversity monitoring and ecological research.

### 2.1 Terminology

The emerging field of automated insect monitoring is generating new concepts and components that require clear and consistent terminology. Inconsistent naming across research groups has created challenges in comparing monitoring systems and protocols. To promote standardization, we propose a set of terms to describe key aspects of hardware design, data collection, and data analysis. Where possible, our terminology is aligned with existing standards for automated biodiversity data collection (Bubnicki et al., 2024; Wieczorek et al., 2012). We use *deployment* to describe a discrete sampling interval associated with one location and one device; *sourceImage* to describe the whole un-cropped image taken by an automated insect monitoring device; and *detection* to refer to a specific subset of a sourceImage that covers only a portion of the sourceImage that contains one item or organism (see Table S3 for a list of terminology).

### 2.2 Mothbox

The Mothbox is an automated light trap for insect monitoring designed to produce accurate insect censuses, while maintaining affordability, durability and portability (Fig. 1). Two Mothbox versions are currently available: Mothbox DIY, which emphasizes device adaptability and supports do-it-yourself (DIY) construction, and Mothbox Pro, which is designed for streamlined manufacturing and large-scale deployment. Full descriptions of components and construction for both Mothbox versions is fully open source with a CERN-OHL-W license (Digital Naturalism Laboratories, 2025a), the operating firmware has an MIT licence, and the documentation is CC BY 4.0. Full specifications, including instructions for construction, programming, use, and adaptation of the Mothbox are on Github (*Mothbox Github*, 2024), and on the Mothbox website (https://mothbox.org/).

**Figure 1.**
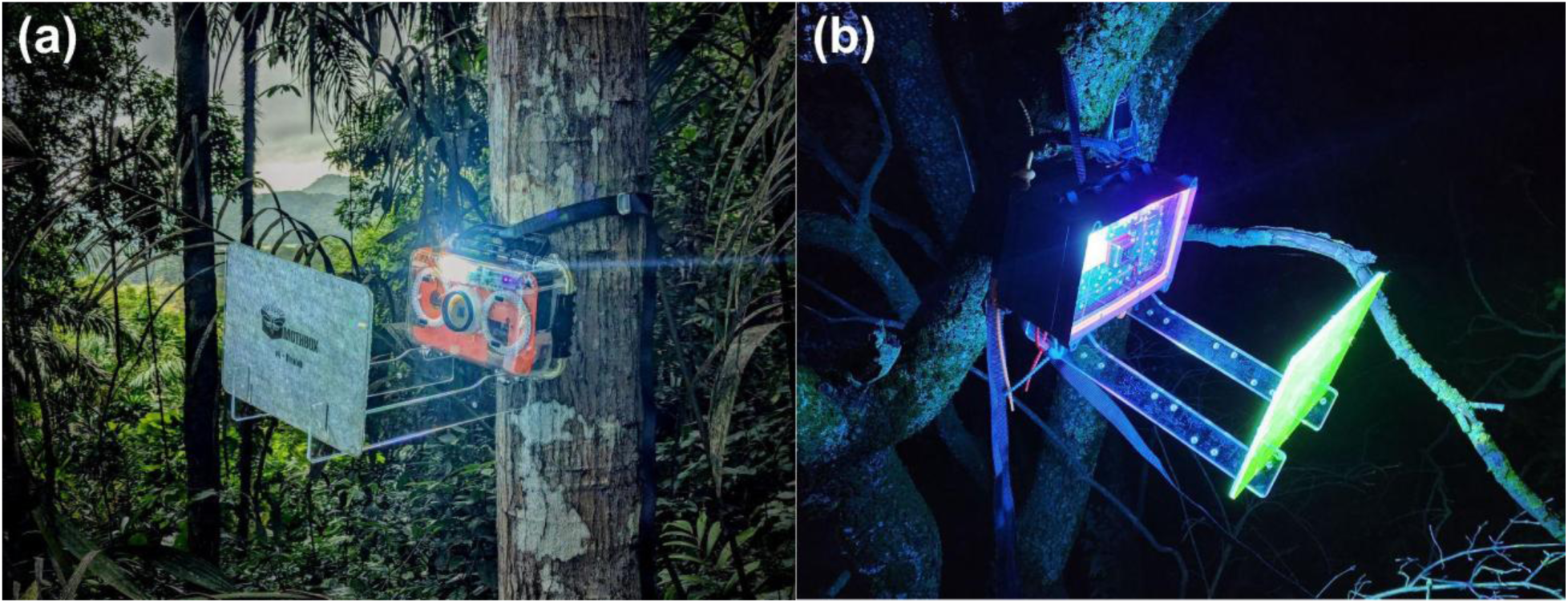
Mothbox DIY (a) and Mothbox Pro (b) deployed in a tropical forest in Panama.

The Mothbox consists of a Raspberry Pi 5 computer that controls a 64 megapixel Arducam Owlsight camera (9248 × 6944 resolution), custom ‘Mothbeam’ lights to attract nocturnal insects, and lights to illuminate the insects for photography (like a camera flash) (Fig. 2). The system is housed in a watertight plastic box and powered by a 156Wh lithium battery. All electronic components are within the watertight housing, however ports in the base of the housing allow for connection to a solar panel, additional batteries, or external light sources. Mothbox Pro models also include an ambient light sensor, internal temperature sensor, slot for attachment of an optional external temperature sensor, and slots for adding an optional GPS unit. Both Mothbox DIY and Mothbox Pro can be programmed via editing a CSV file on the SD card (by plugging the SD card into a different computer or accessing it remotely via a Virtual Network Computing (VNC) system. The Pro model also comes with the addition of manual switches for deployment schedule setting as a backup for easily adjusting settings in the field (Fig. 2). The camera faces a target panel; a 29 cm x 20.5 cm x 0.3 cm acrylic sheet with felted acrylic fabric on one side to create a textured surface diffusely illuminated by the attractant light. The target panel is attached to the Mothbox using clear acrylic arms positioned 28.5cm from the camera. This gives a functional resolution of approximately 625 px / mm^2^ per photograph. At this resolution most insects > 3mm in length can be identified to family level (Fig. 3). The Mothbox can be adapted to use larger target sheets at greater distances from the cameras, which will result in functionally lower resolution photographs (fewer pixels per insect). Mothbox can also be positioned to focus on any flat surface as the camera auto-focuses. Mothbox Pro dimensions are 25.5 x 18 x 13 cm, the arms are 39 x 7 cm, and a full kit with spare arms and a strap weighs 2.7 kg

**Figure 2.**
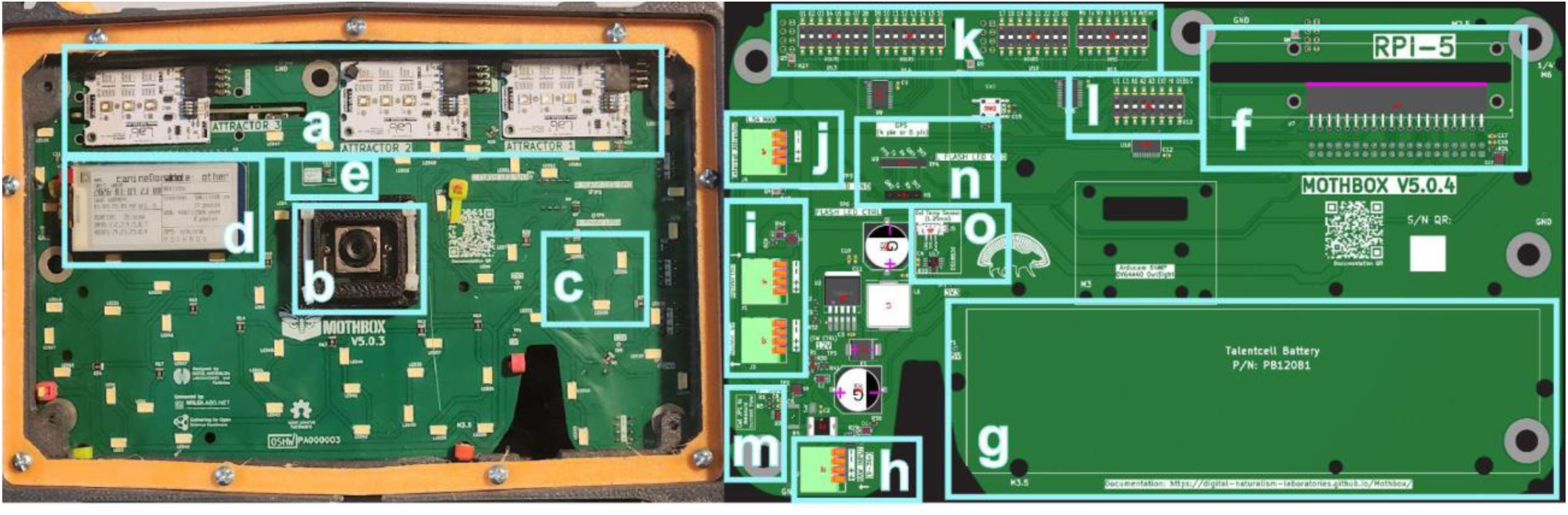
Front (left) and back (right) of Mothbox Pro printed circuit board (PCB) with selected components in blue boxes. The front view is from a complete Mothbox looking in through the clear acrylic front, the back view is a computer-aided design. **a** three Mothbeam PCBs in the attractant light slots; **b** 64 MP Arducam camera; **c** example of three LEDs for illuminating photos (54 total across the front of the PCB); **d** e-paper low-power display with device name, battery status, photo status, and deployment settings; **e** ambient light sensor; **f** Raspberry Pi 5 slot (real time clock battery module attaches to Pi); **g** battery location (voltage regulator attaches to battery, which in turn is fastened to the PCB with zip ties); **h** power raw input (9-36V); **i** 12 V regulator control; **j** external attractant slot; **k** physical schedule programming switches (as a backup option for computer schedule programming); **l** mode programming switches; **m** integrated power monitor; **n** optional 4 or 5 pin GPS slot; **o** temperature sensor with optional external temperature sensor slot.

**Figure 3.**
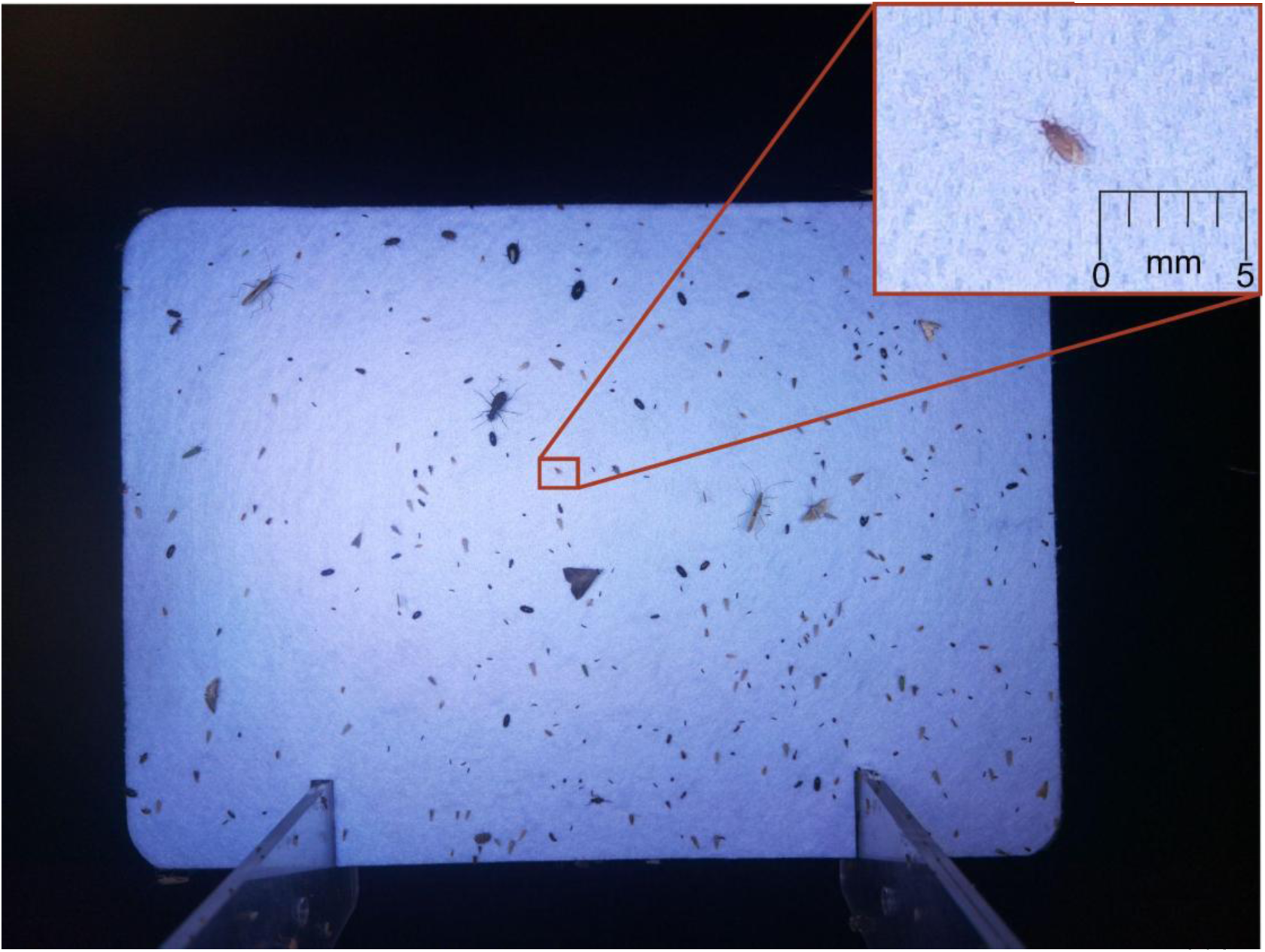
Unedited photo taken by a Mothbox in Panama, with an inset to demonstrate the capacity for documenting small insects like this 1.5 mm long minute pirate bug (Anthocoridae). The resolution of the full photo is 9248 x 6944 pixels. The scale provided is in millimetres.

We have prioritised ease of Do-It-Yourself construction and repair in the Mothbox DIY design. Minimal tools are required for construction (drill, screw driver, wire strippers, epoxy, and a 53mm hole saw drill bit), while most repairs can be managed with a screwdriver, knife, and a hot glue stick. We use lever nuts for electrical connections to avoid soldering to enable modification and repair. The internal framework, arms and target panel can be cut from a single sheet of 50 x 60 x 0.3 cm acrylic. While still 3D-printable, this flat-pack design choice enables easy field servicing, as replacement components can be cut and drilled by hand from acrylic or wood. To set up a Mothbox requires loading a ready-to-go operating system image onto an SD card; no coding expertise is needed. Users can change the sampling schedule by editing a scheduling CSV file in the data storage drive. Mothbox Pro combines the electronic components of Mothbox DIY into a single printed circuit board (PCB), which decreases construction time, number of components, and cost. Mothbox Pro electronics are housed in a custom 3D-printed box, while DIY are in a Plano 1460-00 clear acrylic box. Motion-activated photo saving was the default on the original automated light trap prototype (Bjerge et al., 2021) however for the Mothbox we use time-lapse (photos taken at regular intervals). Time-lapse imaging reduces computational load (and thus extending battery life and eliminating the need for higher-cost Raspberry Pi models), however it can also miss rapid insect activity between frames, potentially underestimating abundance relative to high-sensitivity motion-detection systems. Mothboxes can be programmed for any sampling schedule with no hardware modification. The current default sampling schedule which we use in the tropics is five one-hour intervals at 19:00, 21:00, 23:00, 02:00, and 04:00, with photographs captured every minute (300 images; ∼5.2 GB per night). This intermittent design samples insect assemblages throughout the night, reduces battery consumption and allows the target panel to clear between sessions, promoting insect turnover and reducing repeated observations of the same individuals. Mothbox DIY lasts for 10-12 hours of runtime per battery, or two nights with the default schedule, while Mothbox Pro can last for up to 18 hours, depending on light intensity settings. The single integrated battery can be augmented with an additional box containing up to three more batteries connected in parallel, increasing total run time. A second Plano box with a charging port added to the bottom works for this purpose. The Mothbox can also be connected to portable solar panels (up to 100 watt, 12-20 volt output) and mains power via charging ports in the bottom.

The relatively small size and durability enables the Mothbox to be used in a range of habitats and locations. Once the arms and target panel are in place, and the Mothbox is switched on, it can be mounted to a tree using a strap through the handle (Fig 1), mounted to tripods with the addition of a tripod mount, positioned in the canopy using a throw line, or placed on the ground. Deploying a Mothbox on the ground reduces the range of the attractant lights and gives access to predatory ants and spiders, so elevating the device is advised. After collection, photos are retrieved via a USB drive or SD card and processed with Mothbot.

### 2.3 Mothbeam

The Mothbeam is an open-source printed circuit board (PCB) with light-emitting diodes (LEDs) that emit a range of light frequencies to attract nocturnal insects. The Mothbeam, originally created by Moritz Buttlar of LabLab (Buttlar, 2024), was designed in collaboration with the Mothbox team as an inexpensive, open-source equivalent of a LepiLED (Brehm, 2017). There are two Mothbeam configurations, one which targets violet light (365, 395, 405 nm) and another that targets visible light (full-spectrum white, 450, 520 nm). Mothbeams have four switches that control light intensity. No switches flipped produces the least powerful output (10 mA), two switches add 70mA each, and two switches add 150 mA each. We currently do not advise having all switches engaged at the same time as this might result in overheating the Mothbeam. By default, each Mothbox 4.5 comes with one of each PCB and both PCBs are activated when collecting data. Mothbox 5 has three LED PCB slots. Mothboxes can be programmed to only use one PCB, and the PCBs can be modified to deactivate individual LEDs or dim light intensity, to target particular taxonomic groups or reduce battery consumption. We have also designed stand-alone Mothbeams for use with traditional moth sheeting. Like the Mothbox, Mothbeams are fully open source with a CERN-OHL-W licence (Digital Naturalism Laboratories, 2025b).

### 2.4 Mothbot

Mothbot is an open-source machine learning tool and user interface for processing data from automated light traps. Mothbot consists of two user interfaces: ‘Mothbot Process’ and ‘Mothbot Classify’. The first automatically detects insects from source images, identifies them to a user-defined taxonomic threshold, then spatially clusters them. The second facilitates rapid human validation of computer vision identification and manual finer-taxonomy classification, as well as data export. Mothbot Process uses a combination of two open-sourced AI programs: YOLO (Redmon et al., 2016) and BioCLIP (Stevens et al., 2024). A custom YOLO11 model trained on Mothbox data from 4500 hand-labelled images automatically detects and crops individual insects (down to ∼1 mm total length) from novel imagery. These cropped detections are then fed to a modified BioCLIP model. The model outputs are restricted to taxa on a user-defined species list, generated with taxonomic and geographic filters on GBIF. A custom algorithm then clusters the detections, extracting the detections’ visual features with DINOv2 (https://github.com/facebookresearch/dinov2) and then grouping the detections by similarity with HDBSCAN (Hierarchical Density-Based Spatial Clustering of Applications with Noise - https://pypi.org/project/hdbscan/). These visual clusters are sub-grouped temporally (i.e. by the time their source image was taken) to create hierarchical groupings of detected organisms that exist in temporal ‘tracks’ over a set of sequentially-taken images.

Processing data with Mothbot Process starts with selecting file paths to a folder containing Mothbox JPG photos, a CSV file that contains deployment metadata, and a CSV file containing a species list. The formatting for the metadata CSV file is based on Camtrap DP (Bubnicki et al., 2024) and Darwin Core (Wieczorek et al., 2012) (Table S1). The species list CSV file is in the Global Biodiversity Information Facility (GBIF) downloads format (Table S2), and requires cleaning to remove synonyms and other errors (SI Script 1). For our proof-of-concept studies, our species list included all insect species recorded on GBIF in Panama and Costa Rica. We tried running Mothbot with a species list that also included all arachnids, however, we found that model accuracy decreased with this addition, and data validation was more time-efficient when manually correcting arachnid identifications. After selecting the required file paths, the photos can be processed using the tabs on in the Mothbot Process user interface. The first tab finds detections on the source image, the second identifies them, the third adds metadata to JSON files associated with each sourceImage, the fourth clusters detections perceptually, and the fifth adds EXIF data, including GPS coordinates and identification, to cropped detection photos.

Mothbot Classify is a React-based user interface is designed for human validation of the BioCLIP identifications. Mothbot Classify includes an easily searchable, hierarchical taxonomic classification system, and allows designation of morphospecies that can be linked to iNaturalist observations. The interface exports occurrence data for each organism photographed to a Darwin Core formatted CSV file (Wieczorek et al., 2012). The human-in-the-loop workflow for refining taxonomy is not required for generating an output CSV file, but we believe it is currently an essential step in generating high-quality data from automated insect monitoring devices, as it is for mammal camera trap data (Huebner et al., 2024; Leorna & Brinkman, 2022).

## 3. Proof of concept: Field deployment in Panama

To demonstrate the capabilities of Mothbox devices in field conditions, we conducted proof-of-concept studies examining shifts in insect activity and richness across intra-night temporal dynamics at a single location, and 2. different habitat types assessed simultaneously, to assess whether patterns could be captured through automated sampling with Mothbox hardware and Mothbot processing.

Mothbox development has focused on creating devices that can be deployed in off-grid, remote tropical locations, as no other currently available automated light traps are suited for this use case. The tropics pose some of the most challenging environmental conditions for electronic equipment, including high humidity, elevated temperatures, and biological agents (e.g. insects, rodents and fungi) capable of infiltrating or degrading devices, making tropical forests optimal testing environments for Mothbox robustness and durability. In April 2025, we tested five Mothbox DIY units in two study locations:

1. We deployed one Mothbox with an additional battery for four continuous nights (10 - 13 April 2025) in a tropical forest in the southern Azuero Peninsula, Panama to assess intra-night temporal shifts in insect assemblages.
2. We deployed three Mothboxes simultaneously in a teak plantation, a regenerating forest stand and a neighbouring mature forest near Venao in the southern Azuero Peninsula, Panama for two consecutive nights (11 - 13 April 2025) to assess shifts in insect assemblages with habitat type. Mothbox sampling points were separated by at least 250m.

For both studies we used Mothbox DIY devices mounted on trees at chest height, programmed with a sampling schedule of one-hour sampling sessions starting at 19:00, 21:00, 23:00, 02:00, and 04:00. The Mothboxes were programmed to take a photo every minute, and thus generate 300 images per night. After collecting the Mothboxes from the field, we processed the data using the Mothbot interface set to order-level automatic identification, then manually identified each insect detected to the finest taxonomic level possible and assigned morphospecies where necessary.

## 4 Performance

### 4.1 Detection of patterns in insect number and diversity

In the first study, we recorded 6,271 detections from nine orders, 43 families, and 219 unique morphospecies to assess intra-night temporal shifts in insect assemblages. Only 37 of the 219 morphospecies could be identified to species level by a general entomologist (not an expert taxonomist), the remaining 183 (84%) were assigned morphospecies names. A total of 1046 images were collected over 3 nights (i.e. 19 hours; no data collected 04:00-05:00 on night 4 due to early battery discharge. The target panel had no insects at the beginning of each hour-long session (Fig. 4 c and d), confirming that when the Mothbeam lights were off, all insects left the target panel encouraging insect turnover and reducing repeat recordings. Most insects attracted to the Mothbeams fly around the Mothbox for several minutes before they settle, as they do on conventional moth sheets. During each 1-hr session, the number of detections per photo (Fig. 4 c) and morphospecies richness (Fig. 4 d) increased with time since attractant lights were turned on. The earliest sampling session (19:00-20:00) had the most cumulative detections and the highest morphospecies richness compared to later sampling sessions (Fig. 4 a and b; *P* =0.0002 and 0.0021, respectively, ANOVA).

**Figure 4.**
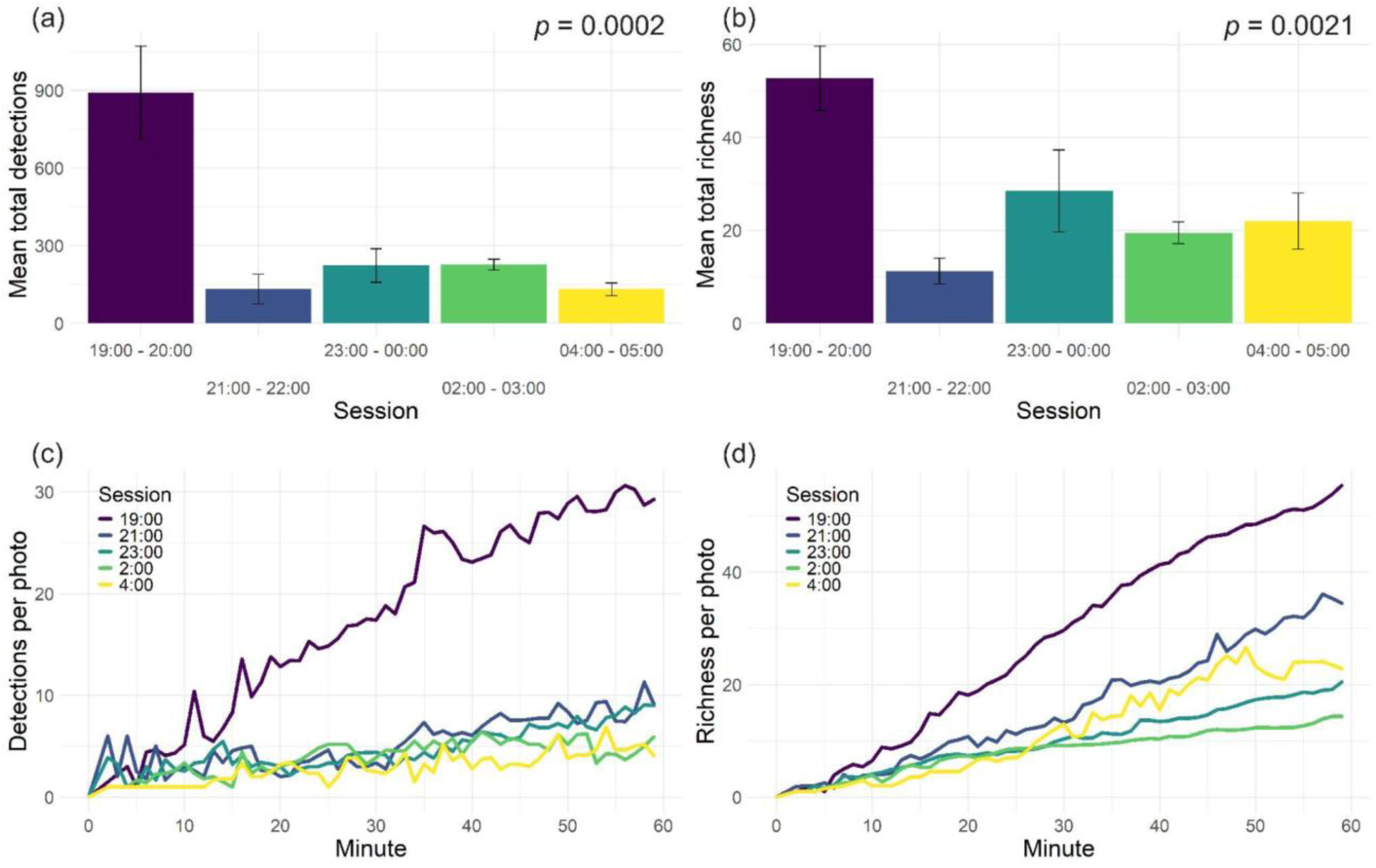
Insect activity proof-of-concept study: (a) total number of detections (individual organisms in each photo, for all photos) for five one-hour sampling sessions spaced throughout the night; (b) total species richness within the five sessions; (c) total number of insect detections in each photo by minute from start of sampling, for the five sampling sessions; (d) species richness per photo across the five sampling sessions. All four panels represent data averaged across the four nights of the deployment. The effect of session on cumulative detections and richness was significant (*p* = 0.0002 and 0.0021, respectively). Error bars in (a) and (b) represent standard error. This study demonstrates that insect activity and richness exhibit intra-night temporal dynamics detectable with automated monitoring

In the second study, we recorded 2,755 detections in total with 8 orders and 40 families represented, and a total of 158 unique morphospecies (with 94% designated morphospecies) across four sampling sites to quantify shifts in insect numbers and species richness with habitat type. Mature forest and native restoration had higher numbers of detections and morphospecies richness (1058 detections / 81 morphospecies and 1140 detections / 64 morphospecies, respectively) than teak plantation (138 detections and 19 morphospecies). Mature forests had higher species richness than the native restoration forest, while native restoration had higher numbers of detections (Fig. 5). Overall, our results illustrate that the Mothbox and Mothbot can effectively detect differences in insect activity and richness across temporal and land-use gradients.

**Figure 5.**
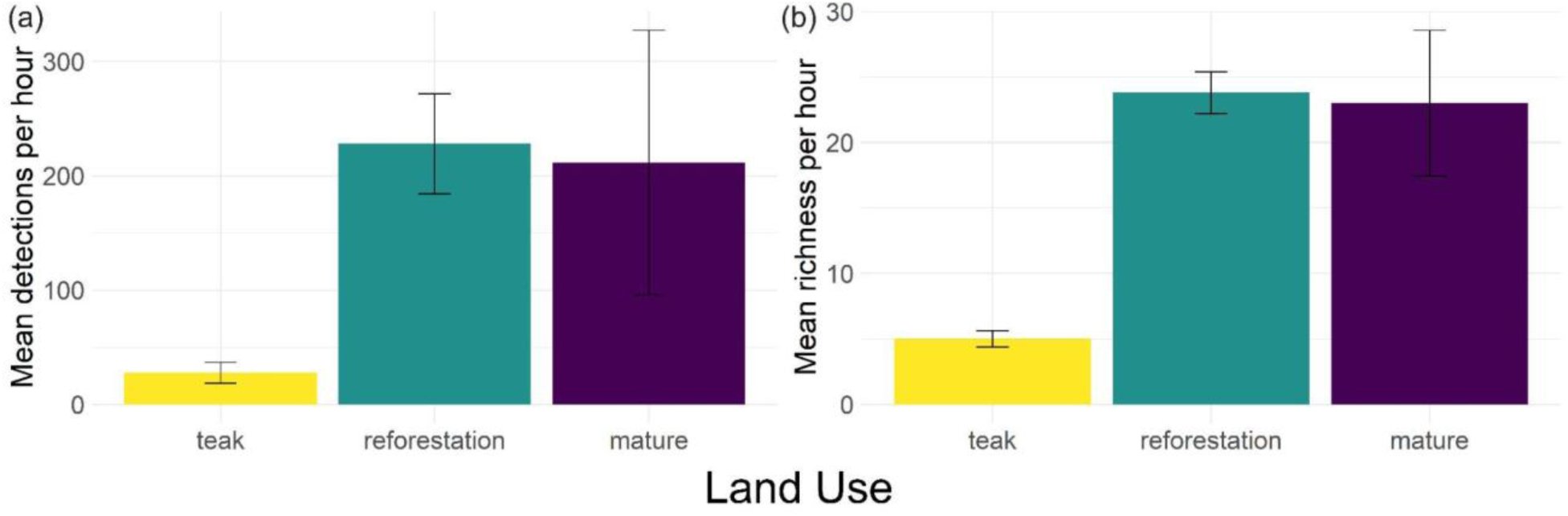
Land use proof-of-concept study. Three Mothboxes were deployed simultaneously for two nights at points with different land use histories. One point was in non-native teak (Tectona grandis) plantation, one point was in an area planted with native tree species, and one point in mature native forest. (a) shows mean detections per hour for each land use type, and (b) shows mean species richness per hour for each land use type.

### 4.2 Identification rates among taxonomic groups

Large, distinctively coloured insects which do not need dissection or microscopic examination were more likely to be identifiable at finer taxonomic levels. For example, of the species which were identified to species level, 87.5% were Lepidoptera, mostly in families Crambidae, Erebidae, and Noctuidae. Conversely, none of the 297 Diptera detections were identified to species level, as most of them were quite small. Some insect detections had poor image resolution due to their small size, and consequently could not be confidently identified to even Order level. For example, it can be difficult to differentiate between the tiniest Diptera and Hymenoptera, or Coleoptera and Hemiptera. While not employed for this analysis, Mothbot Classify has the ability to set size filters, which could be used to narrow the scope of a study to monitoring only larger taxa.

## 5 Implications and Future Directions

### 5.1 Mothbox configuration and considerations

Mothbox can be used to monitor nocturnal insect activity, species richness and composition and can be adapted for other scientific uses. For example, Mothboxes can be modified to release pheromones or nectar from an artificial flower using the spare plug already built-in (Fig. 2 j). If continuous power is available, a Mothbox unit can run on continuous schedules and operate for entire field seasons rather than just short deployments. The Mothbox functions as a standalone sensor, but integrates well with other sensors. For example, acoustic monitors, conventional camera traps, and environmental sensors (light, wind, humidity, precipitation) can all synchronize with Mothbox data collection. This would help identify the drivers of changes in insect activity throughout the night and or across seasons.

While Mothbox development has focused on deployment in remote tropical locations, the system can be applied in any regions or ecosystem where there is an interest in monitoring nocturnal insects. Inclusion of Mothboxes in large-scale ecological monitoring networks like the Long Term Ecological Research (LTER) network and the National Ecological Observatory Network (NEON) would yield an additional valuable dataset for insect population trends (Crossley et al., 2020; Dantzer et al., 2023). Mothbox would also be well suited to monitor pollinator activity and diversity in agriculture, or the spread of invasive species.

### 5.2 Data processing and Mothbot development

The Mothbot Process interface allows users to select the taxonomic level for AI-based identification. Model accuracy depends on the quality and quantity of reference data (in this case training image datasets) and the presence of cryptic species, with finer taxonomic resolution reducing accuracy and increasing manual validation time. In the context of insect identification models, accuracy refers to the correctness of the automated identification relative to a human validation. In our proof-of-concept studies, the model was sufficiently accurate at the order level for rapid human verification, while species-level identification remained unreliable in highly diverse tropical regions with limited reference data. As more detections are manually identified and incorporated into training datasets, future models are expected to achieve reliable family-level identification for most groups and genus- or species-level identification for selected taxa. Our data analysis framework assumes that automated insect monitoring systems support, rather than replace, taxonomic experts, who remain essential for model training and systematic results validation. We therefore prioritized taxonomic accuracy and ease of human verification over fine-scale taxonomic resolution. Where expert taxonomists are unavailable, we recommend uploading representative images of each morphospecies from each deployment to iNaturalist for community-based verification (Campbell et al., 2023).

The accuracy of computer vision models improves when the training data closely matches the data being classified, and Mothbox raw images provide ideal conditions: subjects against a well-lit flat white background, at a fixed distance from the camera. The same cannot be said for the massive and visually diverse datasets that underpin BioCLIP and other current insect classifiers, which include many dead pinned and living but *in-situ* insects. In time, with larger Mothbox datasets, we will augment the Mothbot system with a new model based on human-validated data identified to a finer taxonomic resolution, producing region-specific versions that can accurately identify taxa below family level. We believe that this approach will rapidly change our ability to classify species so that all insects photographed with automated insect monitors can be confidently identified to Genus or Family level by Mothbot. A centralised repository for human-validated image storage and model development in the realm of automated insect monitoring would greatly aid the process of refining classification models, as it has for mammal camera trapping (Vélez et al., 2023). In the meantime, higher taxonomic levels can be used as surrogates of species-level data (Zou et al., 2020), an approach that is key to current large-scale insect monitoring in the tropics while taxonomic expertise, genetic reference libraries, and photographic reference libraries catch up with monitoring needs.

## 5 Conclusion

Mothbox aims to offer for insect monitoring what low-cost camera trapping offers for mammals. By decreasing the time, cost, and specialised knowledge required for monitoring insects, Mothbox enables standardised insect monitoring at the landscape scale. Keeping taxonomists in the loop is critical for ensuring data quality given constraints of insect classification models, and Mothbot offers a platform for rapid validation of computer vision outputs. This technology increases capacity for monitoring global trends in insect populations, and acts as a tool for exploring new ecological relationships and behavioural patterns. Pairing Mothbox and Mothbot has the potential to drive development of human-validated insect image repositories to support improved classification models. Importantly, Mothbox will make biodiversity monitoring more accessible to community conservation groups, empowering restoration and conservation.

## Author contributions

H.A.S and AQ conceived the ideas and designed methodology; H.A.S, B.J., K.Q. and AQ collected the data; H.A.S analyzed the data; H.A.S led the writing of the manuscript; All authors contributed critically to the drafts and gave final approval for publication.

## Acknowledgments

The Mothbox project would not have been possible without many contributors over the years. Special thank you to Wildlabs, Earthshot, Ponterra, Michigan State University, Experiment.com and a Google Carbon Removal Research Award for funding the development of the Mothbox. Thank you to Yash Sondhi for support on early Mothbox development. Thank you to Pro Eco Azuero, Totumas, Daniel Murcia, Josue Santos, Mulget Amaru, and Evidelio Garcia for helping us deploy Mothboxes across Panama. Thank you to the Automated Insect Monitoring community for helping us define terminology.

