## Supplemental information for "Mothbox and Mothbot: automated light trap and data processing system for scalable insect monitoring"

Supporting Information

MothBox: inexpensive, lightweight, automated light trap for scalable insect biodiversity monitoring

**Mothbox website:** https://mothbox.org
**Mothbox github:** https://github.com/Digital-Naturalism-Laboratories/Mothbox/

**Table S1. Sample metadata table**. A CSV file in this format is an input Mothbot. Please see attached CSV file for example.

**Table S2. Species list sample.** A species list that includes all insects catalogued in Panama and Costa Rica on GBIF. Note that this is a tab-delimited text files with UTF-8 encoding and quoted fields, and opening/saving it in Excel will change the format. Please see attached CSV file.

**SI Script 1. R script for cleaning GBIF insect species list CSV files**. This script ensures a unique row for each species, genus, family, and order, removes unaccepted taxa.

**Table S3. Automated Insect Monitoring Terminology.** The set of terminology outlined here assumes stationary devices that capture images.

| **hardware** | |
|---|---|
| device | A machine that performs some kind of automated observation task |
| target | The flat surface being photographed in the sourceImage |
| attractor | The attractant light used. Can also apply to other visual or chemical cues used to attract organisms. |
| attractorDirection | If the attractor is directional, where is it locationed? (for example at the target). If the attractor is not directional 'omnidirectional' can be put here. |
| attractorLocation | Where is the attractor in relation to the camera and the target? For example, 'above target', 'above and between target and device', 'within device', 'within target'. |
| **deployments** | |
| project | a data collection unit composed of several deployments across different locations, sites, and locations |
| site | name for the area being monitored by one or more devices across different sampling locations |
| location | name of a specific sampling location associated with GPS coordinates |

| | |
|---|---|
| | where a device was deployed |
| deployment | name of a 'deployment' - a sampling interval associated with one device at one location, with a single deployentDate and a single collectDate. If two devices are deployed simultaneously at the same location in the same time interval, they will nonetheless be considered to be in two deployments |
| session | Sampling interval of continous monitoring (attractors, if they are present, are activated, images are being captured) |
| sampling schedule | The programmed schedule for a deployment. This will include start and end times of sessions, the timing of sourceImage captures if running the device in timelapse mode, for example. |
| deploymentDate | The start date of the deployment, when the device was deployed at the sampling location (ISO 8601: YYYY-MM-DDThh:mm:ss) |
| collectDate | The end date of the deployment, when the device was collected or moved from the sampling location (ISO 8601: YYYY-MM-DDThh:mm:ss) |
| habitat | a description of the physical environment at the location (radius 50m), for example 'pasture' or 'mature moist tropical forest'. |
| **images and data** | |
| sourceImage | A whole, un-cropped image taken by an automated insect monitoring device, associated with a specific localTime |
| detection | Specific subset of a single sourceImage that covers only a portion of the sourceImage that contains one item (a organism or ERROR) |
| patch | An image file cropped out of a sourceImage corresponding to only one detection |
| track | A series of continuous, sequential detections following one organism |
| organism | The subject of a detection that is an animal (can be an arthropod, a gecko, |
| ERROR | The subject of a detection that is not an animal (dust, a leaf, or moth meconium, for example) |
| classification | Taxonomic name assigned to a detection, either by a human or AI algorithm |
| classifiedBy | Name or identifier of the person or AI algorithm that (most recently) classified the observation |
| occurrenceID | Globally unique name for a detection that depicts one organism |
| scientificName | The full scientific name (for example a latin binomial for organisms identified to species level) |

| | |
|---|---|
| localTime | The time that a sourceImage was taken in the timezone of the location (ISO 8601: YYYY-MM-DDThh:mm:ss) |
| UTC | The time that a sourceImage was taken in UTC (ISO 8601: YYYY-MM-DDThh:mm:ss) |
| attributes | A JSON file with characteristics in a given sourceImage. Its name will be '[sourceImage name]_attributes' |

**Table S4. Parts list for Mothbox Pro.** Prices corrected for one Mothbox, rounded up to the nearest USD.

| Mothbox Pro Parts List | | |
|---|---|---|
| item | estimated price | link |
| **electronics** | | |
| Mothbox mainboard (PCB) | $50 | Instructions: https://digital-naturalism-laboratories.github.io/Mothbox/docs/building/mothbox_pro/manufacture/ |
| Mothbeam boards (up to 3) | $5 | Instructions: https://digital-naturalism-laboratories.github.io/Mothbox/docs/building/mothbox_pro/manufacture/ |
| Raspberry Pi 5 4GB (or higher) | $90 | https://www.sparkfun.com/raspberry-pi-5-4gb.html |
| microSD card (32GB+, Class A1 or faster) | $16 | https://www.amazon.com/dp/B0CQX2B25W?ref=ppx_yo2ov_dt_b_fed_asin_title |
| USB storage (32GB+) | $8 | https://www.amazon.com/dp/B0BN3Q1M96?ref=ppx_yo2ov_dt_b_fed_asin_title |
| Pi 5 RTC battery | $8 | https://www.sparkfun.com/raspberry-pi-rtc-battery.html |
| Talentcell PB120B1 (12V; 9-36V compatible) | $90 | https://www.walmart.com/ip/Millertech-PB120B1-12V-Lithium-ion-Talentcell-Battery-Pack/5119418367 |
| 9-36V to 12V regulator | $19 | https://www.amazon.com/dp/B0B6VK8BPN?ref=ppx_yo2ov_dt_b_fed_asin_title |
| Arducam 64MP Owlsight camera | $60 | https://www.amazon.com/dp/B0CQJPKFVF?ref=ppx_yo2ov_dt_b_fed_asin_title |
| Barrel jack sockets and plugs (x2) | $2 | https://www.amazon.com/dp/B0D9B7WR23?ref=ppx_yo2ov_dt_b_fed_asin_title |
| **3D print/laser cut materials** | | |

| | | |
|---|---|---|
| 3D printed parts (download) | | https://github.com/Digital-Naturalism-Laboratories/Mothbox_Hardware/tree/main/Mothbox_Pro/ |
| Laser-cut parts (download) | | https://github.com/Digital-Naturalism-Laboratories/Mothbox_Hardware/tree/main/Mothbox_Pro/Laser%20Cutting |
| PETG carbon fiber | | https://www.amazon.com/dp/B0FC6H71B1?ref=ppx_yo2ov_dt_b_fed_asin_title |
| PETG (regular) | | https://www.amazon.com/dp/B0FC6H7FBL?ref=ppx_yo2ov_dt_b_fed_asin_title |
| TPU | | https://www.amazon.com/dp/B0B6FMVJ2H?ref=ppx_yo2ov_dt_b_fed_asin_title&th=1 |
| **Other** | | |
| Lever nut connectors | $1 | https://www.amazon.com/Connectors-HTCELLE-60-Piece-Electrical-Terminals/dp/B0C3RQ82BK/?th=1 |
| #6 x 3/8 inch screws | $1 | https://digital-naturalism-laboratories.github.io/Mothbox/docs/building/mothbox_pro/assembleincase/ |
| Thin zip ties (x7) | $1 | https://www.amazon.com/dp/B07W98YW6J?ref=ppx_yo2ov_dt_b_fed_asin_title |
| White felt acrylic (min. ~8x12 in) | $1 | https://www.amazon.com/Jtnohx-Sheets-Handicraft-Fabrics-Projects/dp/B0B493XSSK/?th=1 |
| **Optional** | | |
| Lock (example) | $13 | https://www.amazon.com/gp/product/B000XTPNZK/ref=ppx_yo_dt_b_search_asin_title?ie=UTF8&psc=1 |
| Tree strap | $4 | https://www.amazon.com/gp/product/B08BYHZGC4/ref=ppx_yo_dt_b_search_asin_title?ie=UTF8&psc=1 |
| E-paper (Waveshare 2.13 inch HAT) | $18 | https://www.amazon.com/dp/B0D22JJ18B?ref=ppx_yo2ov_dt_b_fed_asin_title |
| GPS module (5-pin example) | $9 | https://www.amazon.com/dp/B0B31NRSD2?ref_=ppx_hzsearch_conn_dt_b_fed_asin_title_1 |
| GPS module (4-pin example) | $9 | https://www.amazon.com/dp/B0CWL774NR?ref_=ppx_hzsearch_conn_dt_b_fed_asin_title_1 |

**Table S5. Parts list for Mothbox DIY.** Prices corrected for one Mothbox, rounded up to the nearest USD.

| **Mothbox DIY Parts List** | | |
|---|---|---|
| item | estimated price | link |

| electronics | | |
| --- | --- | --- |
| Mothbeam boards (up to 3) | $5 | Instructions: https://digital-naturalism-laboratories.github.io/Mothbox/docs/building/mothbox_pro/manufacture/ |
| Raspberry Pi 5 4GB (or higher) | $90 | https://www.sparkfun.com/raspberry-pi-5-4gb.html |
| microSD card (32GB+, Class A1 or faster) | $16 | https://www.amazon.com/dp/B0CQX2B25W?ref=ppx_yo2ov_dt_b_fed_asin_title |
| USB storage (32GB+) | $8 | https://www.amazon.com/dp/B0BN3Q1M96?ref=ppx_yo2ov_dt_b_fed_asin_title |
| Pi 5 RTC battery | $8 | https://www.sparkfun.com/raspberry-pi-rtc-battery.html |
| Talentcell PB120B1 (12V; 9-36V compatible) | $90 | https://www.walmart.com/ip/Millertech-PB120B1-12V-Lithium-ion-Talentcell-Battery-Pack/5119418367 |
| 9-36V to 12V regulator | $19 | https://www.amazon.com/dp/B0B6VK8BPN?ref=ppx_yo2ov_dt_b_fed_asin_title |
| Arducam 64MP Owlsight camera | $60 | https://www.amazon.com/dp/B0CQJPKFVF?ref=ppx_yo2ov_dt_b_fed_asin_title |
| Barrel jack sockets and plugs (x2) | $2 | https://www.amazon.com/dp/B0D9B7WR23?ref=ppx_yo2ov_dt_b_fed_asin_title |
| USB A to C power cables (short) | $2 | https://www.amazon.com/gp/product/B0C4NJYPTX/ref=ppx_yo_dt_b_search_asin_title?ie=UTF8&th=1 |
| 12V 144-LED ring light (x2 recommended) | $29 | https://www.amazon.com/Vision-Scientific-VMLIFR-09B-Adjustable-Microscope/dp/B07VR2LJJL/ |
| Waveshare 3-channel relay Pi HAT | $18 | https://www.amazon.com/RPi-Relay-Board-Raspberry-3-CH/dp/B085QJFWBC/ |
| 22 AWG solid-core wire (red/black) | $1 | https://www.amazon.com/gp/product/B07JNB712X/ref=ppx_yo_dt_b_search_asin_title?ie=UTF8&th=1 |
| **Other** | | |
| Plano 1460-00 waterproof case | $25 | https://www.amazon.com/gp/product/B003FYMVXM/ref=ppx_yo_dt_b_search_asin_title?ie=UTF8&th=1 |
| UV lens protector / mount rings | $3 | https://www.amazon.com/dp/B005IMNI4K?psc=1&ref=ppx_yo2ov_dt_b_product_details |
| Tiffen 55 mm UV filter | $8 | https://www.amazon.com/dp/B00004ZCJH?psc=1&ref=ppx_yo2ov_dt_b_product_details |
| Lens hood (incl. cap) | $9 | https://www.amazon.com/gp/product/B082HRGFP7/ref=ppx_yo_dt_b_search_asin_title?ie=UTF8&th=1 |

| | | |
|---|---|---|
| M2 / M2.5 / M3 standoffs and screws | $1 | https://www.amazon.com/HVAZI-240pcs-Standoffs-Assortment-Male-Female/dp/B07JYSFMRY/ref=sr_1_2 |
| Lever nut connectors | $1 | https://www.amazon.com/Connectors-HTCELLE-60-Piece-Electrical-Terminals/dp/B0C3RQ82BK/?th=1 |
| #6 x 3/8 inch screws | $1 | https://digital-naturalism-laboratories.github.io/Mothbox/docs/building/mothbox_pro/assembleincase/ |
| Gorilla clear epoxy (plastic) | $2 | https://www.amazon.com/Gorilla-Epoxy-Minute-ounce-Syringe/dp/B001Z3C3AG/ref=sr_1_1 |
| White felt acrylic (min. ~8x12 in) | $1 | https://www.amazon.com/Jtnohx-Sheets-Handicraft-Fabrics-Projects/dp/B0B493XSSK/?th=1 |
| DIY heatsink: perforated aluminum | $6 | https://www.amazon.com/dp/B0CN4RMFYQ/ref=dp_iou_view_item?ie=UTF8&th=1 |
| Thermal silicone strips | $1 | https://www.amazon.com/gp/product/B094PWW9TM/ref=ppx_yo_dt_b_search_asin_title?ie=UTF8&th=1 |
| **Special tools** | | |
| 53 mm hole saw bit | $12 | https://www.amazon.com/Hurricane-02-024-Aviation-Straight-Vanadium/dp/B07GDDF5JB/ref=sr_1_1 |
| Metal shears (example) | $14 | https://www.amazon.com/dp/B07R5RZNQK?psc=1&ref=ppx_yo2ov_dt_b_product_details |
| Stepped / tapered drill bits for plastic | $19 | https://www.amazon.com/Driak-16-30-5-Titanium-Umbrella-Chamfering/dp/B07RX7GRHG/ref=sr_1_3 |
| **Optional** | | |
| Lock (example) | $13 | https://www.amazon.com/gp/product/B000XTPNZK/ref=ppx_yo_dt_b_search_asin_title?ie=UTF8&psc=1 |
| Tree strap | $4 | https://www.amazon.com/gp/product/B08BYHZGC4/ref=ppx_yo_dt_b_search_asin_title?ie=UTF8&psc=1 |
| E-paper (Waveshare 2.13 inch HAT) | $18 | https://www.amazon.com/dp/B0D22JJ18B?ref=ppx_yo2ov_dt_b_fed_asin_title |
| Dielectric grease | $1 | https://www.amazon.com/Mission-Automotive-Dielectric-Silicone-Waterproof/dp/B016E5E59G/ |
